# The Neural Basis of Motivational Influences on Cognitive Control

**DOI:** 10.1101/113126

**Authors:** Cameron Parro, Matthew L Dixon, Kalina Christoff

## Abstract

Cognitive control mechanisms support the deliberate regulation of thought and behavior based on current goals. Recent work suggests that motivational incentives improve cognitive control, and has begun to elucidate the brain regions that may support this effect. Here, we conducted a quantitative meta-analysis of neuroimaging studies of motivated cognitive control using activation likelihood estimation (ALE) and Neurosynth in order to delineate the brain regions that are consistently activated across studies. The analysis included functional neuroimaging studies that investigated changes in brain activation during cognitive control tasks when reward incentives were present versus absent. The ALE analysis revealed consistent recruitment in regions associated with the frontoparietal control network including the inferior frontal sulcus (IFS) and intraparietal sulcus (IPS), as well as consistent recruitment in regions associated with the salience network including the anterior insula and anterior mid-cingulate cortex (aMCC). A large-scale exploratory meta-analysis using Neurosynth replicated the ALE results, and also identified the caudate nucleus, nucleus accumbens, medial thalamus, inferior frontal junction/premotor cortex (IFJ/PMC), and hippocampus. Finally, we conducted separate ALE analyses to compare recruitment during cue and target periods, which tap into proactive engagement of rule-outcome associations, and the mobilization of appropriate viscero-motor states to execute a response, respectively. We found that largely distinct sets of brain regions are recruited during cue and target periods. Altogether, these findings suggest that flexible interactions between frontoparietal, salience, and dopaminergic midbrain-striatal networks may allow control demands to be precisely tailored based on expected value.

## Introduction

The ability to maintain attention during a lecture, or flexibly shift between writing a report and answering emails, or plan several steps ahead during a chess match all require cognitive control—the capacity to deliberately guide thought and behavior based on goals, especially in the presence of distraction or competing responses (Botvinick et al., 2001; Desimone & Duncan, 1995; Duncan, 2013; Gollwitzer, 1999; Miller & Cohen, 2001; Miyake et al., 2000; Posner & Dehaene, 1994; Posner & DiGirolamo, 1998; Stuss & Knight, 2002). Cognitive control involves several related, yet dissociable abilities (Miyake et al., 2000), including working memory (D'Esposito & Postle, 2015; Funahashi, Chafee, & Goldman-Rakic, 1993; Fuster & Alexander, 1971; Goldman-Rakic, 1987), representation of rules and context (Asaad, Rainer, & Miller, 2000; Bunge, 2004; Cohen & Servan-Schreiber, 1992; Dixon & Christoff, 2012; Koechlin, Ody, & Kouneiher, 2003; Miller & Cohen, 2001; Munakata et al., 2011), conflict and error detection (Botvinick et al., 2001; Ridderinkhof, Ullsperger, Crone, & Nieuwenhuis, 2004; Ullsperger, Danielmeier, & Jocham, 2014), inhibition of pre-potent responses (Aron, Robbins, & Poldrack, 2004), abstract thought and reasoning (Christoff et al., 2009; Christoff et al., 2001; Dias, Robbins, & Roberts, 1996), and set-shifting (Crone, Wendelken, Donohue, & Bunge, 2006; Meiran, 1996; Meiran, 2000; Rushworth, Passingham, & Nobre, 2002).

While early work identified the prefrontal cortex (PFC) as a critical neural substrate (Desimone & Duncan, 1995; Duncan, 2001; Fuster, 1989; Miller & Cohen, 2001; Passingham & Wise, 2012; Stuss & Knight, 2002; Watanabe, 2017), it soon became clear that a much broader network of regions support cognitive control, including posterior parietal, lateral temporal, insular, and mid-cingulate cortices, as well as parts of the basal ganglia. Together, these regions are often referred to as the frontoparietal control network (FPCN) or Multiple Demand system (Cole, Repovs, & Anticevic, 2014; Cole et al., 2013; Cole & Schneider, 2007; Crittenden, Mitchell, & Duncan, 2016; Dixon, Andrews-Hanna, et al., 2017; Dixon, Girn, & Christoff, 2017; Dosenbach et al., 2007; Duncan, 2010; Mitchell et al., 2016; Spreng et al., 2010; Vincent et al., 2008). The FPCN flexibly represents a variety of task-relevant information, and exerts a top-down influence on other regions, guiding activation in accordance with current task demands (Buschman & Miller, 2007; Crowe et al., 2013; Desimone & Duncan, 1995; Dixon, Fox, & Christoff, 2014b; Egner & Hirsch, 2005; Miller & Cohen, 2001; Tomita et al., 1999).

### The effects of motivation on cognitive control

As research progressed in delineating the components of cognitive control, a separate stream of inquiry focused on the neural mechanisms of assigning value to stimuli and value-guided decision making (Daw, Niv, & Dayan, 2005; Dixon & Christoff, 2014; Dixon, Thiruchselvam, Todd, & Christoff, 2017; Levy & Glimcher, 2012; O'Doherty, 2004; Rangel, Camerer, & Montague, 2008; Rangel & Hare, 2010; Rushworth et al., 2011; Schoenbaum & Esber, 2010). The past decade has seen a synthesis of these fields with a surge of interest in understanding how value influences the decision of whether or not to engage cognitive control and the efficacy of implementing control (Botvinick & Braver, 2015; Braver et al., 2014; Cohen, Braver, & Brown, 2002; Cools, 2016; Dixon, 2015; Dixon & Christoff, 2012; Hazy, Frank, & O'Reilly R, 2007; McGuire & Botvinick, 2010; O'Reilly, Herd, & Pauli, 2010). This line of inquiry is yielding new insights into mechanisms that allow the desire to achieve a specific outcome to interact with the cognitive processes that are necessary to realize that outcome, and may ultimately provide critical information about pathological conditions that involve altered motivation-cognition interactions including depression, schizophrenia, ADHD, and anxiety (Barkley, 1997; Bishop, Duncan, Brett, & Lawrence, 2004; Chung & Barch, 2015; Davidson, 2000; Heller et al., 2009; Kaiser, Andrews-Hanna, Spielberg, et al., 2015; Kaiser, Andrews-Hanna, Wager, & Pizzagalli, 2015; Nigg & Casey, 2005; Pessoa, 2008; Shackman et al., 2011; Shackman et al., 2016).

Recent studies have shown that individuals are strongly biased towards choosing habits and simple tasks over more complex or demanding tasks that require cognitive control (Botvinick & Braver, 2015; Dixon & Christoff, 2012; Kool, McGuire, Rosen, & Botvinick, 2010; McGuire & Botvinick, 2010). This has led to notion that cognitive control carries an intrinsic *effort cost*. This effort cost can be offset by the opportunity to acquire a rewarding outcome. Studies have shown that participants are considerably more likely to engage cognitive control if doing so will result in a larger reward than if they chose a habitual action (Dixon & Christoff, 2012; Westbrook, Kester, & Braver, 2013). Thus, cognitive control engagement can be understood as a special case of cost/benefit decision making whereby the expected value of the outcome that will result from engaging cognitive control is weighed against the effort cost of its implementation (Botvinick & Braver, 2015; Dixon & Christoff, 2012; Shenhav, Botvinick, & Cohen, 2013).

Following the decision to engage cognitive control, the opportunity to earn a reward can also influence the efficacy of implementing control processes. In one study, participants performed a modified Stroop task during which they decided whether an image was a building or a house, and had to ignore letters overlaid on the images (Padmala & Pessoa, 2011). The letters could be neutral (XXXXX), congruent with the image (e.g., HOUSE printed over a house image), or incongruent (e.g., BLDNG printed over a house image). Pre-trial cues indicated whether monetary rewards were available or not available, and participants could only earn rewards if performance was fast and accurate. The results demonstrated enhanced implementation of cognitive control, manifest as reduced interference effects on incongruent trials when rewards were available (Padmala & Pessoa, 2011). This incentive effect may reflect a sharpening of the representation of task-relevant information (Etzel et al., 2015; Histed, Pasupathy, & Miller, 2009), thus providing more effective modulation of sensorimotor processes that support performance. Incentive-based facilitation of behavioral performance has been reported across numerous studies using a range of cognitive control paradigms (Chiew & Braver, 2013, 2014; Chiew, Stanek, & Adcock, 2016; Dixon & Christoff, 2012; Etzel et al., 2015; Ivanov et al., 2012; Jimura, Locke, & Braver, 2010; Krebs et al., 2012; Locke & Braver, 2008; Padmala & Pessoa, 2011; Taylor et al., 2004).

### The neural basis of motivational effects on cognitive control

Functional neuroimaging studies have identified brain regions associated with the influence of motivation on the implementation of cognitive control (Bahlmann, Aarts, & D'Esposito, 2015; Beck et al., 2010; Engelmann, Damaraju, Padmala, & Pessoa, 2009; Gilbert & Fiez, 2004; Ivanov et al., 2012; Kouneiher, Charron, & Koechlin, 2009; Locke & Braver, 2008; Padmala & Pessoa, 2011; Pochon et al., 2002; Rowe, Eckstein, Braver, & Owen, 2008; Taylor et al., 2004). In one study, Jimura and colleagues (2010) employed a Sternberg task with two types of task blocks. One block consisted of only non-reward trials, while the other block consisted of trials with varying outcomes: no reward, low reward ($0.25), or high reward ($0.75). On each trial participants were presented with a 5-word memory set and then had to indicate whether a subsequent probe word matched one of the items in the memory set. The results demonstrated a shift from transient to sustained activation in lateral prefrontal and parietal cortices during reward versus no reward blocks, and individual differences in reward sensitivity correlated with the magnitude of sustained activation in reward contexts (Jimura et al., 2010).

These results can be interpreted in terms of the dual mechanisms of control (DMC) framework, which suggests that reward incentives shift the type and timing of cognitive control (Braver, 2012; Chiew & Braver, 2013; Jimura et al., 2010). This theory posits two temporally-defined cognitive control mechanisms: (i) a *proactive* mechanism consisting of sustained activation of task-relevant information (e.g., task rules) across trials, which facilitates the encoding of new information on each trial and the preparation of a target response; and (ii) a *reactive* mechanism consisting of the stimulus-triggered transient re-activation of rule information on a trial-by-trial basis. Frontoparietal activation dynamics support the idea that reward incentives lead to greater reliance on proactive control, consistent with the DMC model.

Numerous studies have now observed elevated frontoparietal activation when cognitive control is performed in the service of obtaining rewarding outcomes (Boehler et al., 2014; Engelmann et al., 2009; Gilbert & Fiez, 2004; Ivanov et al., 2012; Kouneiher et al., 2009; Locke & Braver, 2008; Padmala & Pessoa, 2011; Paschke et al., 2015; Pochon et al., 2002; Rowe et al., 2008; Soutschek et al., 2015; Taylor et al., 2004). Additionally, frontoparietal regions encode associations between specific rules and expected reward outcomes (Dixon & Christoff, 2012), exhibit more differentiated coding of task rules on incentivized trials (Etzel et al., 2015), and are sensitive to the interaction between control level and reward availability (Bahlmann et al., 2015; Ivanov et al., 2012; Padmala & Pessoa, 2011; Soutschek et al., 2015). These regions are also recruited during value-based decision making, and when participants plan and monitor progress towards future desired outcomes (Crockett et al., 2013; Dixon, Fox, & Christoff, 2014a; Gerlach, Spreng, Madore, & Schacter, 2014; Jimura, Chushak, & Braver, 2013; McClure, Laibson, Loewenstein, & Cohen, 2004). Finally, single cell recordings in non-human primates have revealed reward-contingent enhancement of lateral PFC neural firing related to working memory and task rules (Histed et al., 2009; Leon & Shadlen, 1999; Watanabe, 1996; Watanabe & Sakagami, 2007). Thus, frontoparietal regions may integrate task-relevant information and expected motivational outcomes (Dixon & Christoff, 2014; Pessoa, 2008; Watanabe, 2017; Watanabe & Sakagami, 2007).

### The current meta-analysis

While numerous studies of motivated cognitive control have reported activation in frontoparietal regions, the consistency of activations across these studies has yet to be systematically examined. The present study sough to characterize the network of brain regions that are consistently recruited during motivated cognitive control. To this end we used a quantitative approach, activation likelihood estimation (ALE), to identify regions that show consistent recruitment in human neuroimaging studies of cognitive control that included a manipulation of reward incentive availability. We additionally used Neurosynth to identify regions that are consistently recruited in studies that use the term “cognitive control” *and* in studies that use the term “reward”. While the ALE analysis provides a conservative and rigorous analysis based on a set of carefully selected studies, the Neurosynth analysis provides a complementary perspective based on a liberal exploration of a much wider literature. Finally, we performed two additional exploratory ALE analyses to examine activations during cue and target periods. During cue periods, participants are presented with information about task rules for responding to stimuli and expected payoffs. This period thus allows for preparatory construction rule-outcome associations in service of proactive control engagement. During target periods, participants respond to stimuli and must mobilize appropriate viscero-motor states to facilitate faster and more accurate behaviors when a reward is on the line. This analysis allowed us to examine the extent to which cue and target periods rely on similar versus distinct brain systems.

## Materials and Methods

### Search strategy

We conducted a literature search through PubMed and Google Scholar to identify peer-reviewed neuroimaging studies that have investigated motivated cognitive control. We began by searching the key terms “fMRI” AND (“reward” OR “motivation”) AND (“cognitive control” OR “executive function” OR “working memory”). We then read the abstract of each paper to confirm or reject it as a candidate study for inclusion in the meta-analysis. We only focused on activations, because there are very few deactivations reported in the literature. Additionally, we focused on the effect of reward, because only a few studies have looked at the effect of punishment. To be included in the analysis, studies had to fulfill the following criteria: (i) employ fMRI and report resulting activation coordinates; (ii) include a cognitive control task (e.g., Stroop) with a manipulation of motivational incentive (i.e., reward versus no reward, or high versus low reward conditions); (iii) include healthy adult human participants; and (iv) report results from a whole-brain analysis. Several studies of motivated cognitive control employed ROI-based analyses and were not included in the meta-analysis, given that ALE requires whole-brain analyses to provide unbiased results. Sixteen studies were found that matched the inclusion criteria (**Table 1**). The presence of reward was associated with significantly improved behavioral performance (decreased reaction time and/or increased accuracy) in all but one of the sixteen studies.

**Table 1.**
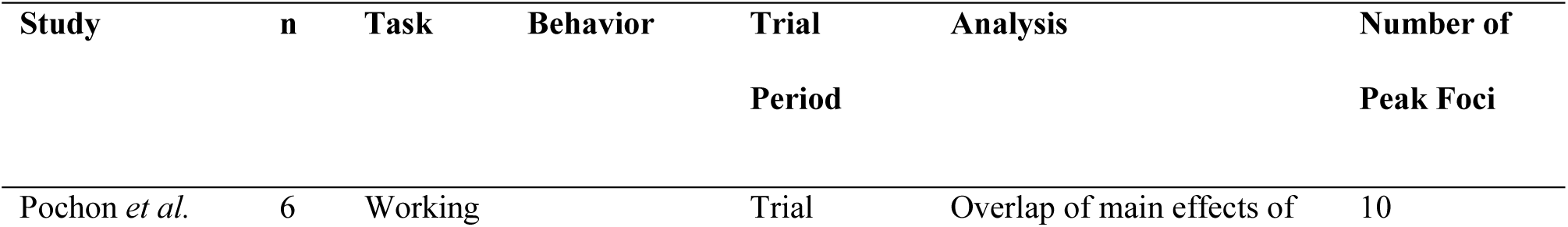
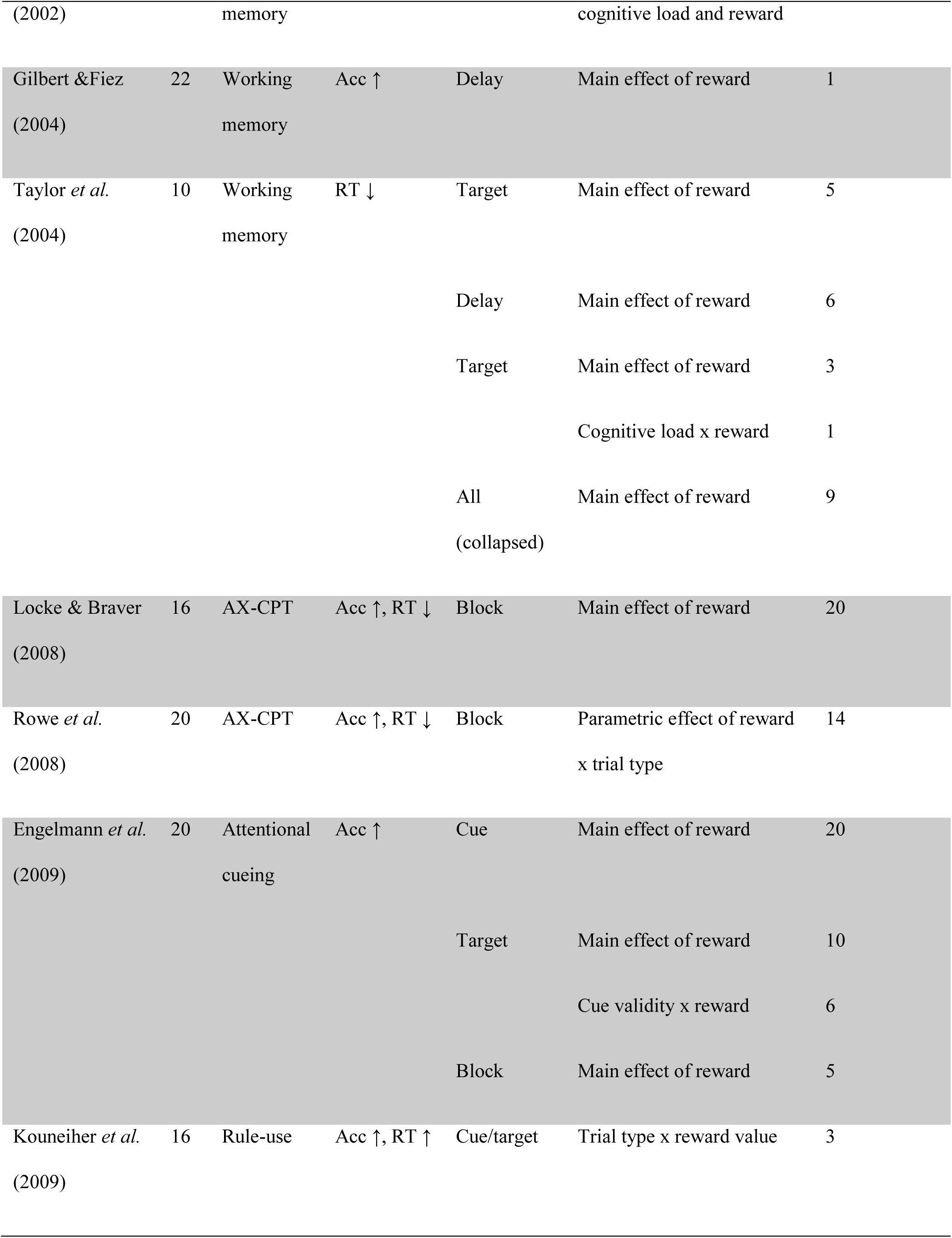
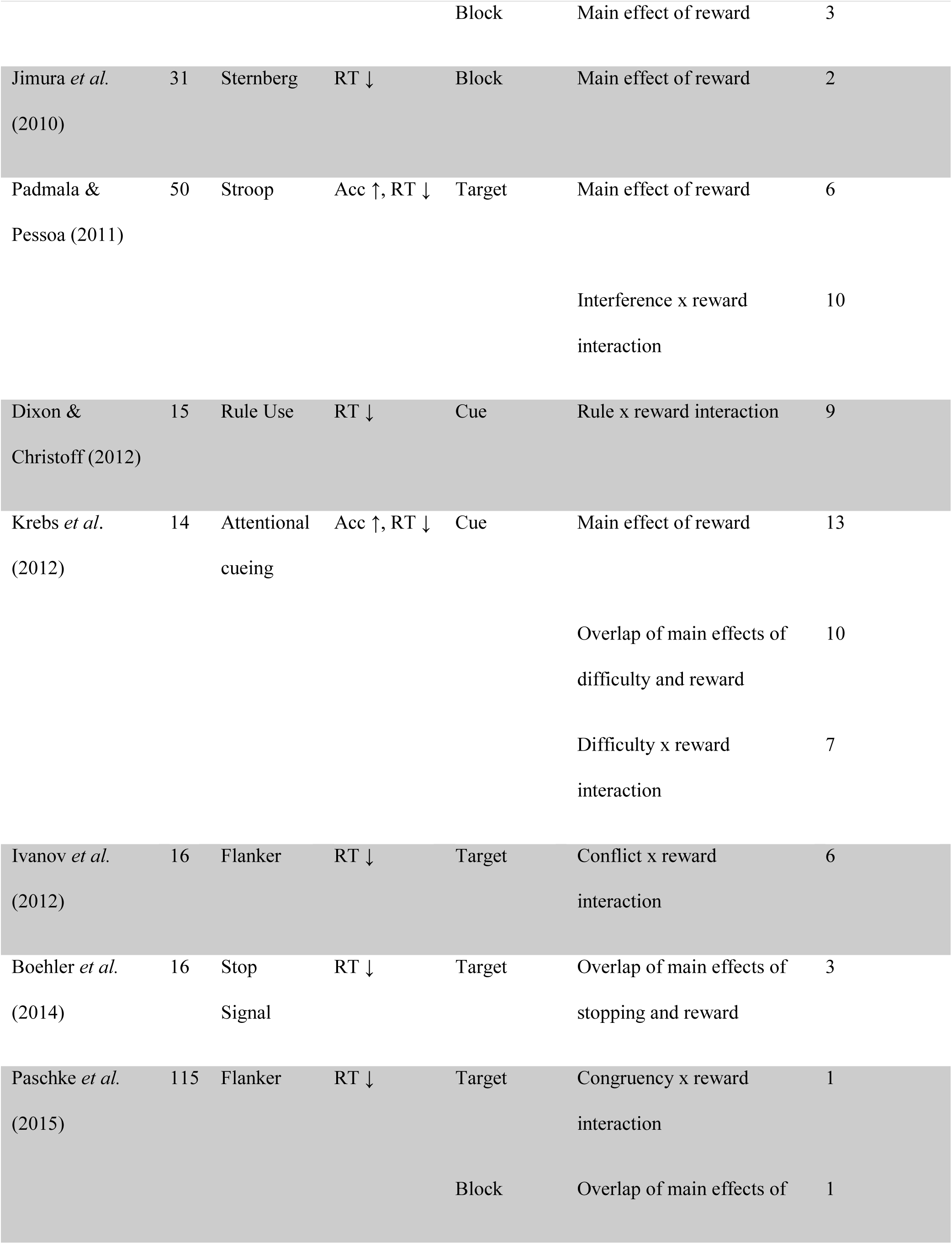
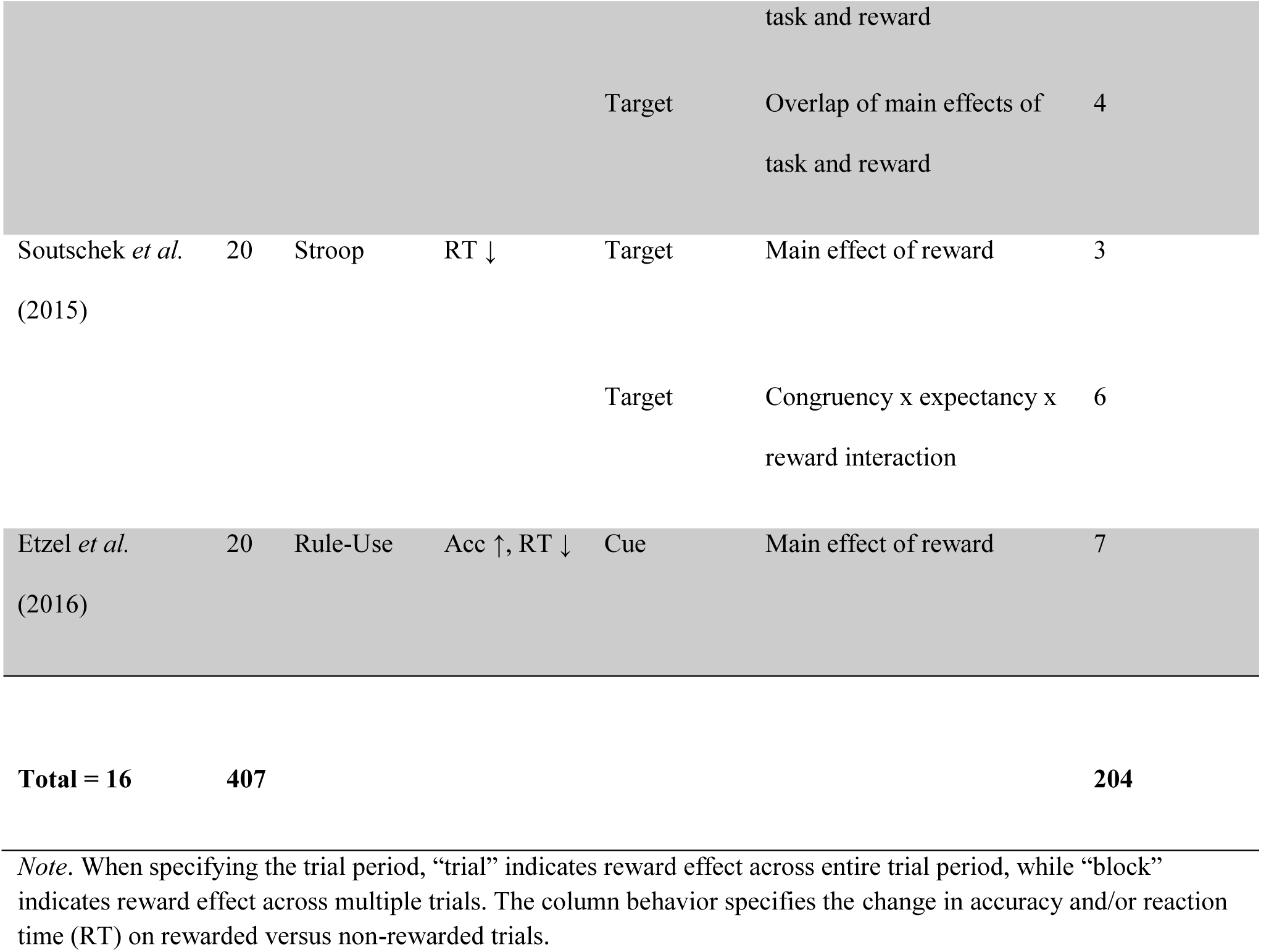
Studies included in the meta-analysis.

### Data extraction

From these sixteen studies, we collected data on sample size, task, type of contrast (e.g., main effect of reward during task, or reward x cognitive load interaction), task period (e.g., cue, delay, or target), and peak activation coordinates (**Table 1**). The meta-analysis included studies with different types of contrasts, but each examined the neural substrates that link motivational incentives to cognitive control. There were three categories of contrasts: (i) main effect of reward during a cognitive control task; (ii) conjunction effects showing overlapping activation in relation to cognitive demands and sensitivity to reward value; and (iii) interaction between cognitive control level and presence of incentive. While there are some differences in these three types of contrasts, all converge on related processes that support incentive-based modulation of cognitive control. It should be noted that we included results from the main effect of reward during task performance (e.g., during delay or target periods) but excluded results related to a main effect of reward during cue periods that *only* revealed the expected reward incentive, as this is likely to mainly capture reward processing alone, without an interaction with cognitive processes. If the cue period signaled motivational information *and* cognitive information (e.g., rules) that could be activated in a preparatory manner, then we included these foci. For studies that had multiple periods (e.g., delay, probe), we included foci from each period; however, if a given brain region was activated in multiple periods, it was only included once in the meta-analysis. Note that for the separate cue period and target period analyses, all available foci were used.

### ALE meta-analytic data analysis

We analyzed the activation coordinates using a random-effects meta-analysis, activation likelihood estimation (ALE) (Eickhoff et al., 2012; Eickhoff et al., 2009; Laird et al., 2005; Turkeltaub et al., 2012) implemented with GingerALE 2.3.6 software (San Antonio, TX: UT Health Science Center Research Imaging Institute). This is the updated version of GingerALE that has fixed the error related to cluster-level FWE correction (Eickhoff et al., 2017).

Coordinates reported in Talairach space were first converted to MNI space using GingerALE’s foci converter function: Talairach to MNI (SPM). ALE models the uncertainty in localization of activation foci across studies using Gaussian probability density distributions. The voxel-wise union of these distributions yields the ALE value, a voxel-wise estimate of the likelihood of activation, given the input data. The algorithm aims at identifying significantly overlapping clusters of activation between studies. ALE treats activation foci from single studies as 3D Gaussian probability distributions to compensate for spatial uncertainty. The width of these distributions was statistically determined based on empirical data for between subject and between template variability (Eickhoff et al., 2009). Additionally, studies were weighted according to sample size, reflecting the idea that large sample sizes more likely reflect a true localization. This is implemented in terms of a widening Gaussian distribution with lower sample sizes and a smaller Gaussian distribution (and thus a stronger impact on ALE scores) with larger sample sizes (Eickhoff et al., 2009). Modeled activation maps for each study were generated by combining the probabilities of all activation foci for each voxel (Turkeltaub et al., 2012). These ALE scores were then compared to an ALE null distribution (Eickhoff et al., 2012) in which the same number of activation foci was randomly relocated and restricted by a gray matter probability map (Evans, Kamber, Collins, & MacDonald, 1994). Spatial associations between experiments were treated as random while the distribution of foci within an experiment was treated as fixed. Thereby random effects inference focuses on significant convergence of foci between studies rather than convergence within one study. The ALE scores from the actual meta-analysis were then tested against the ALE scores obtained under this null-distribution yielding a p-value based on the proportion of equal or higher random values. For the main ALE analysis, we used a cluster-forming threshold at the voxel level of *p* < 0.001, and a cluster-level threshold of *p* < 0.05 FWE corrected for multiple comparisons. We also ran separate analyses on foci from the cue period and foci from the target period. Given that fewer studies were included in each analysis, we used a more liberal *p* < .001, uncorrected threshold, with a minimum cluster size of 200 mm^3^. Results were visualized with MRIcron software (Rorden, Karnath, & Bonilha, 2007).

### Neurosynth meta-analyses

To examine consistent recruitment related to motivated cognitive control using a more liberal and exploratory approach, we performed term-based forward inference meta-analyses using Neurosynth (Yarkoni et al., 2011). To perform such automated meta-analyses, Neurosynth divides the entire database of coordinates into two sets: those that occur in articles containing a particular term, and those that don't. A large-scale meta-analysis is then performed comparing the coordinates reported for studies with and without the term of interest. Forward inference maps reflect z-scores corresponding to the likelihood that each voxel will activate if a study uses a particular term (P(Activation|Term)), and are corrected for multiple comparisons using a false discovery rate (FDR) of *q* = .01. Here, we conducted forward inference meta-analyses using the terms “cognitive control” and “reward”, and looked for brain areas demonstrating overlapping recruitment across both domains.

## Results

### ALE meta-analysis: all foci

We first performed an analysis on all foci to identify regions that consistently demonstrate increased activation during cognitive control when rewards are available versus not available. The ALE analysis revealed four large clusters (**Figures 1 and 2; Table 2**). These right-lateralized regions included the inferior frontal sulcus (IFS) extending into the mid-dorsolateral prefrontal cortex (mid-DLPFC), mid-intraparietal sulcus (mid-IPS) extending into the anterior inferior parietal lobule (aIPL), anterior insula, and the anterior mid-cingulate cortex (aMCC) extending into the adjacent pre-supplementary motor area (pre-SMA).

**Figure 1.**
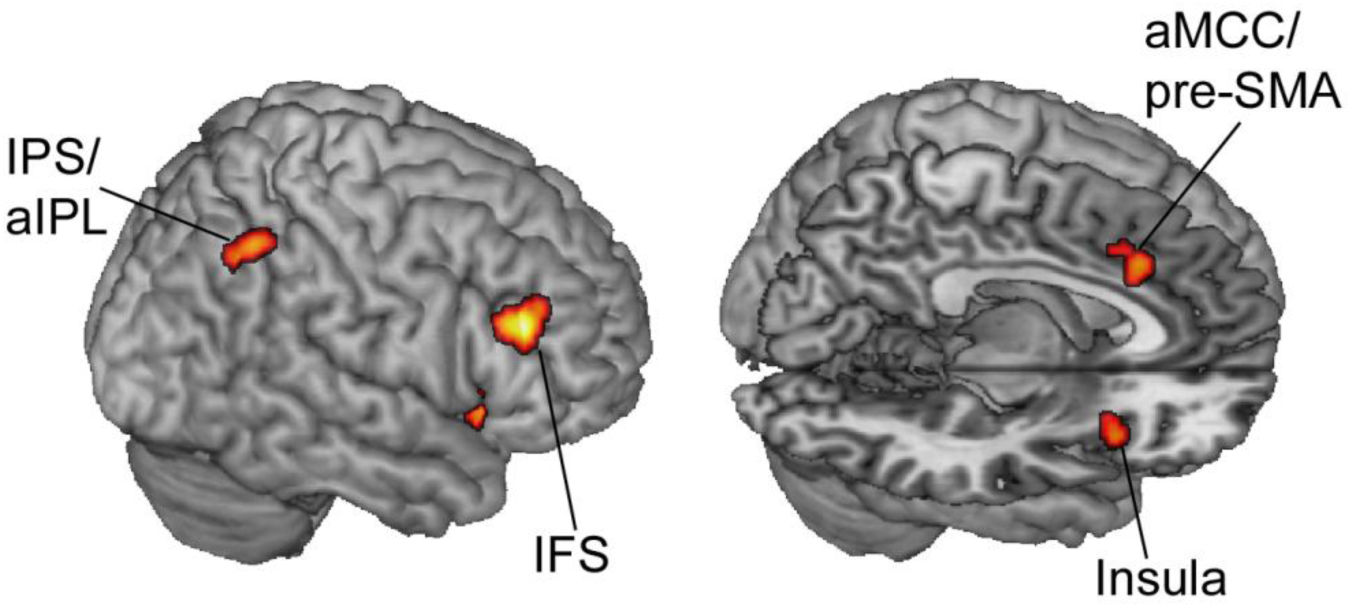
Rendered surface display of significant ALE meta-analytic clusters (*p* < .05 FWE corrected). Abbreviations: IFS, inferior frontal sulcus; IPS/aIPL, intraparietal sulcus/anterior inferior parietal lobule; aMCC/pre-SMA, anterior mid-cingulate cortex/pre-supplementary motor area.

**Figure 2.**
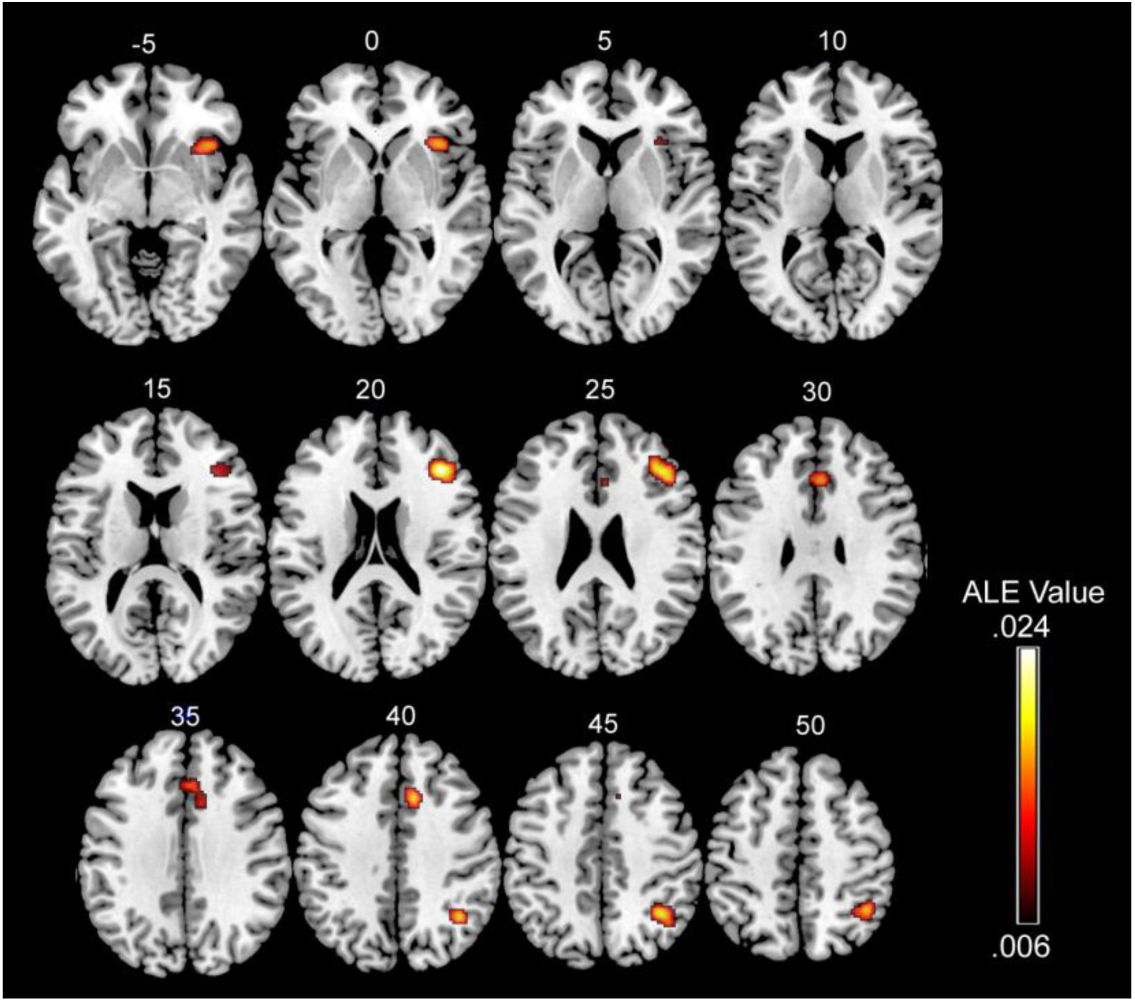
Whole-brain ALE meta-analytic results (*p* < .05 FWE corrected). Numbers denote z-coordinates in MNI space.

**Table 2.** Significant ALE clusters

| Region | Cluster<br>size (mm<sup>3</sup>) | Peak ALE<br>value | Weighted peak foci in<br>MNI space (x,y,z) |
| --- | --- | --- | --- |
| R inferior frontal sulcus / mid-dorsolateral prefrontal cortex | 2032 | 0.0252 | 40, 32, 22 |
| R anterior insula | 1560 | 0.0168 | 38, 20, -2 |
| R intraparietal sulcus / inferior parietal lobule | 1544 | 0.0213 | 39, -53, 45 |
| R anterior mid-cingulate cortex / pre-supplementary motor area | 1288 | 0.0172 | 10, 20, 40 |

*Note.* Abbreviations: L, left; R, right; DLPFC, dorsolateral prefrontal cortex.

### Neurosynth meta-analyses

Although the strict inclusion criteria in the ALE analysis offers confidence that the identified regions do play a key role in motivated cognitive control, it is possible that this conservative analysis may overlook other relevant regions. Thus, as a complementary analysis, we used Neurosynth to identify regions that are consistently activated in studies that use the term “cognitive control” (N = 428 studies) and in studies that use the term “reward” (N = 671 studies). We focused on brain areas demonstrating overlapping recruitment across these domains, and may play a role in linking control demands to motivational outcomes. Notably, the regions demonstrating this pattern included all of the regions identified in the ALE meta-analysis (**Figure 3**). This analysis additionally identified homologous regions in the left hemisphere, as well as the bilateral inferior frontal junction/pre-motor cortex (IFJ/PMC), bilateral caudate nucleus extending into the nucleus accumbens (NAcc), bilateral medial thalamus, and bilateral hippocampus (**Figure 3**).

**Figure 3.**
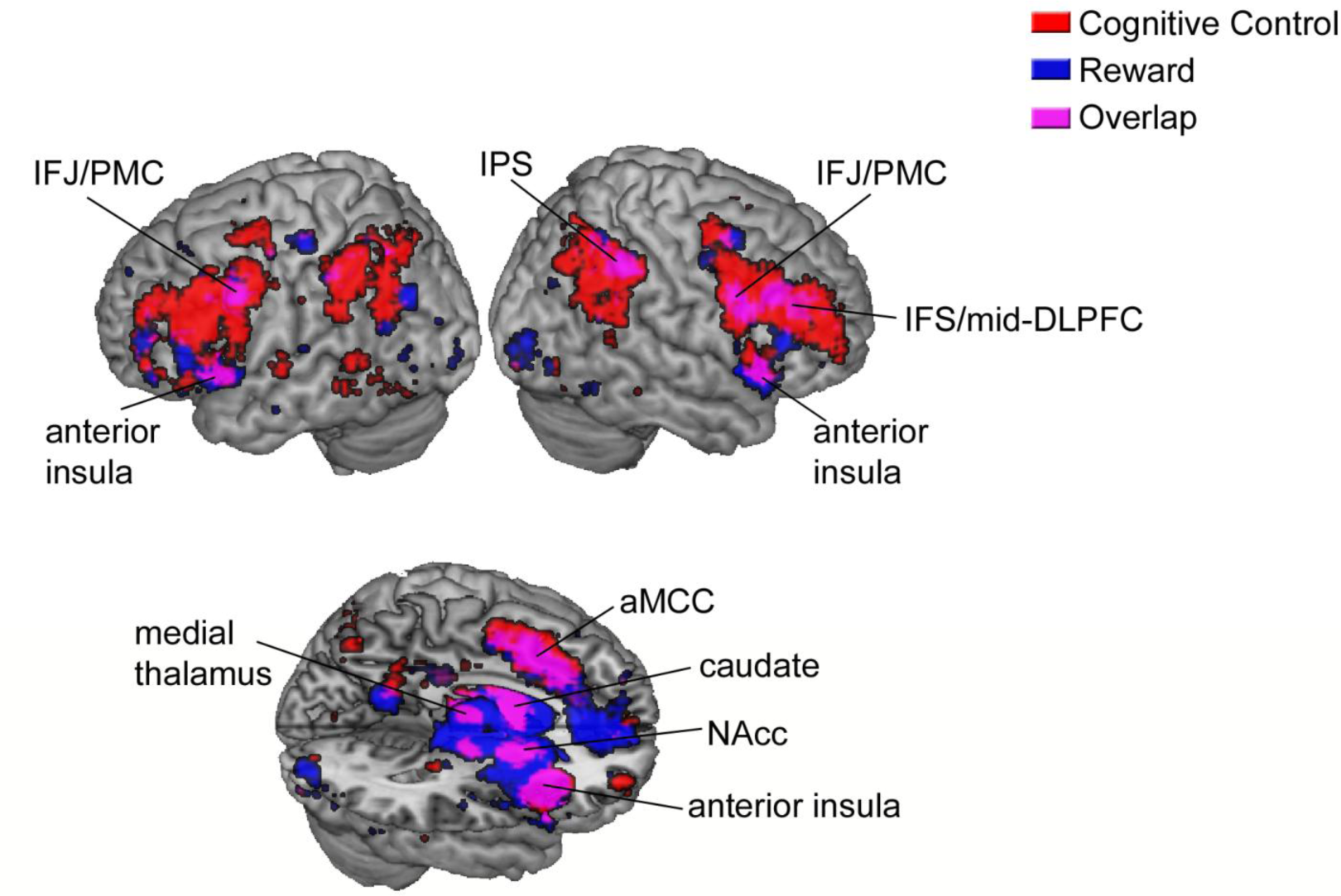
Rendered surface display of Neurosynth forward inference meta-analyses using the terms “cognitive control” and “reward”, corrected for multiple comparisons using a false discovery rate of *q* = .01. Abbreviations: IFS, inferior frontal sulcus; IFJ/PMC, inferior frontal junction/premotor cortex; IPS, intraparietal sulcus; DLPFC, dorsolateral prefrontal cortex; aMCC, anterior mid-cingulate cortex; NAcc, nucleus accumbens.

### ALE meta-analyses: cue and target period foci

In our final analysis, we examined similarities and differences in neural recruitment during cue and target periods. Given that these analyses were based on a limited number of studies and a more lenient statistical threshold, these results should be viewed as exploratory. The brain regions that were consistently recruited during the cue period, and may contribute to the proactive engagement of value-modulated control signals, included the right IFJ/PMC, left ventral IPS, bilateral caudate, right dorsal posterior cingulate cortex (PCC), right midbrain near the ventral tegmental area (VTA), and left medial thalamus (**Figure 4**). On the other hand, the brain regions that were consistently recruited during the target period, and may contribute to the mobilization of viscero-motor processes during action selection, included the right anterior insula, right aMCC, right IPS/aIPL, right medial thalamus, left ventral IPS, and left IFJ (**Figure 4)**. The only region common to both trial events was the left ventral IPS, suggesting that value-based modulation of control processes during cue and target periods may rely on largely distinct neural systems. However, this is a tentative conclusion, tempered by the low power of these analyses.

**Figure 4.**
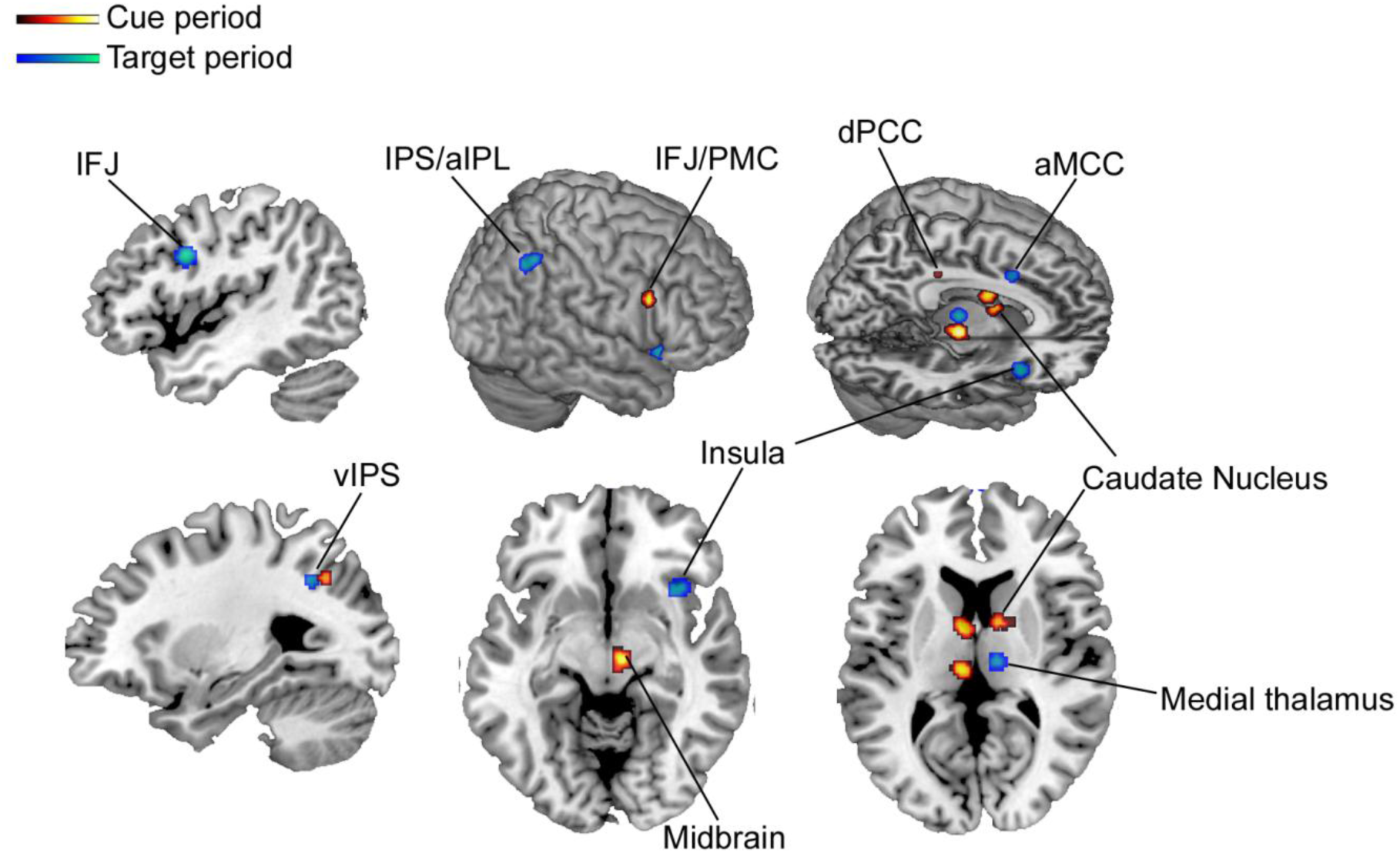
ALE meta-analytic clusters associated with the cue period and the target period (*p* < .001 uncorrected). Abbreviations: IPS/aIPL, intraparietal sulcus/anterior inferior parietal lobule; vIPS, ventral intraparietal sulcus; dPCC, dorsal posterior cingulate cortex; aMCC, anterior mid-cingulate cortex; IFJ, inferior frontal junction; PMC, pre-motor cortex.

## Discussion

Cognitive control is often enhanced when reward incentives are contingent on performance. This enhancement manifests as faster and more accurate responses, and is often accompanied by elevated brain activation in numerous cortical regions. Here, we sought to characterize the brain regions that reliably demonstrate this pattern and may support incentive-related behavioral improvements in cognitive control. Using quantitative ALE and Neurosynth meta-analyses, we identified a select constellation of multimodal association cortices and subcortical regions known to play key roles in motivational processing. An exploratory analysis also revealed differences in recruitment during cue versus target periods, suggesting partially distinct systems may underlie the proactive engagement of control versus the mobilization of viscero-motor states that support action execution.

Several regions were implicated in both the ALE and Neurosynth analyses including the inferior frontal sulcus (IFS), intraparietal sulcus (IPS)/anterior inferior parietal lobule (aIPL), anterior mid-cingulate cortex (aMCC)/pre-supplementary motor area (pre-SMA), and anterior insula. The fact that similar results were obtained with different analysis criteria provides strong evidence that these regions are centrally involved in value-based modulation of cognitive control. Interestingly, the Neurosynth analysis revealed that only a subset of regions engaged during cognitive control are also engaged during reward processing. For example, the posterior middle temporal gyrus and parts of the lateral prefrontal and parietal cortices only demonstrated consistent recruitment during cognitive control. This suggests that there may be a select group of regions including the IFS, IPS/aIPL, aMCC/pre-SMA, and insula that integrate control demands and expected outcomes.

The aforementioned regions have well-established roles in supporting cognitive control and adaptive behavior via top-down modulation of sensory and motor processing (Cole et al., 2014; Dosenbach et al., 2006; Duncan, 2010; Miller & Cohen, 2001). The IFS and IPS/aIPL are part of the frontoparietal control network (FPCN) (Dixon, Girn, et al., 2017; Power et al., 2011; Spreng et al., 2010; Vincent et al., 2008; Yeo et al., 2011) and contribute to working memory and the flexible representation of task rules (Badre & D'Esposito, 2009; Brass, Derrfuss, Forstmann, & von Cramon, 2005; Bunge, 2004; De Baene, Kuhn, & Brass, 2011; Derrfuss, Brass, Neumann, & von Cramon, 2005; Dixon & Christoff, 2012; Dumontheil, Thompson, & Duncan, 2011; Koechlin et al., 2003; Wallis, Anderson, & Miller, 2001). Neurons in these regions exhibit dynamic coding properties, signaling any currently relevant information (Duncan, 2010; Stokes et al., 2013), and rapidly updating their pattern of global functional connectivity according to task demands (Cole et al., 2013; Fornito, Harrison, Zalesky, & Simons, 2012; Gao & Lin, 2012; Spreng et al., 2010). One possibility is that elevated activation during motivated cognitive control reflects an amplification and sharpening of task information (e.g., rules) as a result of modulatory inputs from reward processing regions (Cohen et al., 2002; Etzel et al., 2015; Histed et al., 2009; Kouneiher et al., 2009). It could also reflect a shift in the temporal dynamics of cognitive control, towards a proactive mode of control (Braver, 2012; Jimura et al., 2010). When performance needs to be fast and accurate in order to procure a reward, FPCN regions exhibit greater sustained activation and reduced transient/reactive activation, ostensibly reflecting the active maintenance of task rules across trials (Braver, 2012; Jimura et al., 2010).

Several lines of evidence suggest that FPCN regions may play an integrative role, directly representing control demands in relation to expected outcomes. First, Dixon & Christoff (2012) found that the FPCN flexibly represented trial-to-trial shifts in the association between specific task rules and expected reward outcomes (Dixon & Christoff, 2012). This finding is consistent with the fact that FPCN neurons encode not only rule information, but also experienced and expected reward and punishment (Matsumoto, Suzuki, & Tanaka, 2003; Pan et al., 2008; Wallis & Miller, 2003; Watanabe, Hikosaka, Sakagami, & Shirakawa, 2002)(Abe & Lee, 2011; Asaad & Eskandar, 2011; Hikosaka & Watanabe, 2000; Histed et al., 2009; Hosokawa & Watanabe, 2012; Kennerley & Wallis, 2009; Kim, Hwang, & Lee, 2008; Klein, Deaner, & Platt, 2008; Kobayashi et al., 2006; Platt & Glimcher, 1999; Seo, Barraclough, & Lee, 2007; Watanabe, 1996; Watanabe et al., 2002). Second, McGuire and Botvinick (2010) found that the lateral PFC signaled the cost of exerting cognitive effort, suggesting that the FPCN plays a role in linking control demands to other parameters that are important for deciding when to implement control. In fact, numerous studies have demonstrated FPCN activation during value-based decision making (Christopoulos et al., 2009; Diekhof & Gruber, 2010; Gianotti et al., 2009; Huettel, Song, & McCarthy, 2005; Hutcherson, Plassmann, Gross, & Rangel, 2012; Jimura et al., 2013; Jimura & Poldrack, 2012; Lebreton et al., 2013; McClure et al., 2004; Plassmann, O'Doherty, & Rangel, 2007; Plassmann, O'Doherty, & Rangel, 2010; Tanaka et al., 2004; Tobler et al., 2009; Vickery, Chun, & Lee, 2011; Weber & Huettel, 2008). Third, several studies have shown an interaction between control level and reward expectancy in the FPCN (Bahlmann et al., 2015; Ivanov et al., 2012; Padmala & Pessoa, 2011). Finally, disruption of the FPCN—via transcranial magnetic stimulation or due to a lesion—disrupts value processing and leads to altered motivation (Camus et al., 2009; Essex, Clinton, Wonderley, & Zald, 2012; Paradiso et al., 1999; Zamboni et al., 2008). Together, these findings suggest that the FPCN may play an integrative role, serving as a bridge between control demands and motivational outcomes (Dixon & Christoff, 2014; Dixon, Thiruchselvam, et al., 2017; Pessoa, 2008; Watanabe, 2017; Watanabe & Sakagami, 2007).

Our meta-analytic results also revealed that the anterior insula and anterior mid-cingulate cortex play key roles in motivated cognitive control. These regions are part of the “salience network” (Menon & Uddin, 2010; Seeley et al., 2007). The insula has a well-established role in interoception—the representation of internal bodily signals including pain, temperature, respiratory and cardiac sensations (Craig, 2002; Critchley & Harrison, 2013; Critchley et al., 2004; Farb, Segal, & Anderson, 2012). This region is also activated during a variety of goal-directed tasks (Dixon et al., 2014a; Dosenbach et al., 2006; Duncan, 2010; Farb et al., 2012), suggesting that it may serve as a nexus between the FPCN and other interoceptive regions, allowing viscero-somatic signals to become integrated with task goals (Dixon et al., 2014a; Farb et al., 2012; Jezzini et al., 2012). It has been suggested that the aMCC plays a role in determining the value of implementing control (Shenhav et al., 2013). An alternative perspective is that this region plays a more specific role in liking reinforcement to different actions (Camille, Tsuchida, & Fellows, 2011; Dixon, Thiruchselvam, et al., 2017; Rushworth, Behrens, Rudebeck, & Walton, 2007). This region is well connected to the motor system (Morecraft & Tanji, 2009), is sensitive to the effort costs of actions (Croxson et al., 2009; Kurniawan, Guitart-Masip, Dayan, & Dolan, 2013; Shidara & Richmond, 2002), and encodes action-outcome associations (Alexander & Brown, 2011; Procyk et al., 2014; Rushworth et al., 2007; Shackman et al., 2011). Unlike the IFS and IPS, the aMCC does not encode rule-outcome associations (Dixon & Christoff, 2012). Rather, during motivated cognitive control, the insula and aMCC may play a role in translating rule-outcome associations represented by the FPCN into concrete viscero-motor body states that drive optimal behavior in service of acquiring a reward or avoiding punishment (Dixon et al., 2014a; Knutson & Greer, 2008; Medford & Critchley, 2010; Rushworth et al., 2011; Shima & Tanji, 1998). Consistent with this idea, we found that the anterior insula and aMCC were specifically recruited during the target rather than cue period. The functions of these regions may facilitate the maintenance of effort prior to and during action execution (Croxson et al., 2009; Parvizi et al., 2013; Shidara & Richmond, 2002).

The Neurosynth analysis also highlighted the caudate nucleus extending into the NAcc, while the cue period ALE analysis highlighted the caudate and the midbrain near the VTA. These regions are part of a dopaminergic midbrain-striatal circuit that signals prediction errors when there is a discrepancy between expected and actual rewards (Hare et al., 2008; Montague, Dayan, & Sejnowski, 1996; O'Doherty et al., 2004; Schultz, 1997). Moreover, neurons in this circuit gradually shift the timing of maximal firing from actual outcomes to reward-predictive cues (Schultz, 1997). These regions play a fundamental role in goal-directed behavior via biasing action selection based on the anticipation of imminent rewards and the opportunity to exercise choice (Knutson, Adams, Fong, & Hommer, 2001; Leotti & Delgado, 2014). Accordingly, this circuit may play a role in broadcasting predicted cue values to other systems involved in constructing rule-outcome associations, and modulating viscero-motor processing. Indeed, prior work has outlined detailed models of how the dopaminergic midbrain-striatal circuit serves a gating function that strengthens or destabilizes current working memory contents depending on task demands (Cohen et al., 2002; Cools, 2016; Hazy, Frank, & O'Reilly, 2006). Specifically, tonic dopamine in the PFC is thought to enhance the stability of working memory content via increased signal to noise ratio (that is, boosting the strength of local recurrent activity versus stimulus-evoked activity). On the other hand, phasic dopamine is thought to serve as a gating signal, allowing working memory to be updated based on reward-predicting events (Cohen et al., 2002; Cools, 2016; Hazy et al., 2006).

A few limitations of the current findings should be noted. Our analysis was based on studies that employed a number of different cognitive control tasks. One the one-hand, this suggests that the identified brain regions support motivated cognitive control in general and are not tied to any particular task. On the other hand, this may obscure the delineation of neural systems that may link expected outcomes to different types of executive control. As more studies examine this topic, future work may be able to discern whether incentive effects on different aspects of cognitive control (e.g., response inhibition versus working memory updating) have similar or distinct neural substrates. We were also unable to examine the effect of punishment on cognitive control given the small number of fMRI studies on this topic. Given that the observed frontoparietal regions have been shown to encode information about aversive outcomes in addition to rewarding outcomes (Asaad & Eskandar, 2011; Kobayashi et al., 2006), it is possible that substantial overlap with the current findings would be observed. However, there is some indication in prior work that differences may also appear (Paschke et al., 2015). Future studies may also be able to provide a more in-depth analysis of brain regions showing incentive effects during specific trial periods (e.g., cue versus delay and target processing). Our results were based on a small number of studies and should be seen as preliminary. Given that we found evidence of distinct brain regions involved in cue versus target periods, this may be a key area for future inquiry to investigate. Another important dimension of motivated cognitive control is incentive type (i.e., primary versus secondary). However, all studies included in this review operationalized motivation with monetary (i.e., secondary) incentives with the exception of Beck *et al.* (2010). This study compared the effects of primary (juice) and secondary (money) rewards on performance in a Sternberg task. The authors found no significant differences in behavioral improvement between the reward types, but did find both regional and temporal differences in brain activation patterns. This underscores the importance of studying the different types of incentive effects separately.

To summarize, the current findings reveal a select constellation of brain regions that are consistently recruited in studies of motivated cognitive control. Flexible interactions between frontoparietal, salience, and dopaminergic midbrain-striatal networks may underlie the dynamic process by which control signals are precisely tailored based on expected outcomes.

## References

Abe, H., & Lee, D. (2011). Distributed coding of actual and hypothetical outcomes in the orbital and dorsolateral prefrontal cortex. Neuron, 70(4), 731–741.

Alexander, W. H., & Brown, J. W. (2011). Medial prefrontal cortex as an action-outcome predictor. Nat Neurosci, 14(10), 1338–1344.

Aron, A. R., Robbins, T. W., & Poldrack, R. A. (2004). Inhibition and the right inferior frontal cortex. Trends Cogn Sci, 8(4), 170–177.

Asaad, W. F., & Eskandar, E. N. (2011). Encoding of both positive and negative reward prediction errors by neurons of the primate lateral prefrontal cortex and caudate nucleus. J Neurosci, 31(49), 17772–17787.

Asaad, W. F., Rainer, G., & Miller, E. K. (2000). Task-specific neural activity in the primate prefrontal cortex. Journal of neurophysiology, 84(1), 451–459.

Badre, D., & D’Esposito, M. (2009). Is the rostro-caudal axis of the frontal lobe hierarchical? Nat. Rev. Neurosci, 10(9), 659–669.

Bahlmann, J., Aarts, E., & D'Esposito, M. (2015). Influence of motivation on control hierarchy in the human frontal cortex. J Neurosci, 35(7), 3207–3217.

Barkley, R. A. (1997). Behavioral inhibition, sustained attention, and executive functions: constructing a unifying theory of ADHD. Psychol Bull, 121(1), 65–94.

Beck, S. M., Locke, H. S., Savine, A. C., Jimura, K., & Braver, T. S. (2010). Primary and secondary rewards differentially modulate neural activity dynamics during working memory. PloS one, 5(2), e9251.

Bishop, S., Duncan, J., Brett, M., & Lawrence, A. D. (2004). Prefrontal cortical function and anxiety: controlling attention to threat-related stimuli. Nat Neurosci, 7(2), 184–188.

Boehler, C. N., Schevernels, H., Hopf, J. M., Stoppel, C. M., & Krebs, R. M. (2014). Reward prospect rapidly speeds up response inhibition via reactive control. Cognitive, affective & behavioral neuroscience, 14(2), 593–609.

Botvinick, M., & Braver, T. (2015). Motivation and cognitive control: from behavior to neural mechanism. Annu Rev Psychol, 66, 83–113.

Botvinick, M. M., Braver, T. S., Barch, D. M., Carter, C. S., & Cohen, J. D. (2001). Conflict monitoring and cognitive control. Psychological review, 108(3), 624–652.

Brass, M., Derrfuss, J., Forstmann, B., & von Cramon, D. Y. (2005). The role of the inferior frontal junction area in cognitive control. Trends Cogn Sci, 9(7), 314–316.

Braver, T. S. (2012). The variable nature of cognitive control: a dual mechanisms framework. Trends Cogn Sci, 16(2), 106–113.

Braver, T. S., Krug, M. K., Chiew, K. S., Kool, W., Westbrook, J. A., Clement, N. J., et al. (2014). Mechanisms of motivation-cognition interaction: challenges and opportunities. Cogn Affect Behav Neurosci, 14(2), 443–472.

Bunge, S. A. (2004). How we use rules to select actions: a review of evidence from cognitive neuroscience. Cogn Affect Behav Neurosci, 4(4), 564–579.

Buschman, T. J., & Miller, E. K. (2007). Top-down versus bottom-up control of attention in the prefrontal and posterior parietal cortices. Science, 315(5820), 1860–1862.

Camille, N., Tsuchida, A., & Fellows, L. K. (2011). Double dissociation of stimulus-value and action-value learning in humans with orbitofrontal or anterior cingulate cortex damage. J Neurosci, 31(42), 15048–15052.

Camus, M., Halelamien, N., Plassmann, H., Shimojo, S., O'Doherty, J., Camerer, C., et al. (2009). Repetitive transcranial magnetic stimulation over the right dorsolateral prefrontal cortex decreases valuations during food choices. The European journal of neuroscience, 30(10), 1980–1988.

Chiew, K. S., & Braver, T. S. (2013). Temporal dynamics of motivation-cognitive control interactions revealed by high-resolution pupillometry. Front Psychol, 4, 15.

Chiew, K. S., & Braver, T. S. (2014). Dissociable influences of reward motivation and positive emotion on cognitive control. Cognitive, affective & behavioral neuroscience, 14(2), 509–529.

Chiew, K. S., Stanek, J. K., & Adcock, R. A. (2016). Reward Anticipation Dynamics during Cognitive Control and Episodic Encoding: Implications for Dopamine. Frontiers in human neuroscience, 10, 555.

Christoff, K., Keramatian, K., Gordon, A. M., Smith, R., & Madler, B. (2009). Prefrontal organization of cognitive control according to levels of abstraction. Brain research, 1286, 94–105.

Christoff, K., Prabhakaran, V., Dorfman, J., Zhao, Z., Kroger, J. K., Holyoak, K. J., et al. (2001). Rostrolateral prefrontal cortex involvement in relational integration during reasoning. NeuroImage, 14(5), 1136–1149.

Christopoulos, G. I., Tobler, P. N., Bossaerts, P., Dolan, R. J., & Schultz, W. (2009). Neural correlates of value, risk, and risk aversion contributing to decision making under risk. The Journal of neuroscience : the official journal of the Society for Neuroscience, 29(40), 12574–12583.

Chung, Y. S., & Barch, D. (2015). Anhedonia is associated with reduced incentive cue related activation in the basal ganglia. Cognitive, affective & behavioral neuroscience, 15(4), 749–767.

Cohen, J. D., Braver, T. S., & Brown, J. W. (2002). Computational perspectives on dopamine function in prefrontal cortex. Current opinion in neurobiology, 12(2), 223–229.

Cohen, J. D., & Servan-Schreiber, D. (1992). Context, cortex, and dopamine: a connectionist approach to behavior and biology in schizophrenia. Psychological Review, 99(1), 45-77.

Cole, M. W., Repovs, G., & Anticevic, A. (2014). The frontoparietal control system: a central role in mental health. The Neuroscientist : a review journal bringing neurobiology, neurology and psychiatry, 20(6), 652–664.

Cole, M. W., Reynolds, J. R., Power, J. D., Repovs, G., Anticevic, A., & Braver, T. S. (2013). Multi-task connectivity reveals flexible hubs for adaptive task control. Nat Neurosci, 16(9), 1348–1355.

Cole, M. W., & Schneider, W. (2007). The cognitive control network: Integrated cortical regions with dissociable functions. Neuroimage, 37(1), 343–360.

Cools, R. (2016). The costs and benefits of brain dopamine for cognitive control. Wiley interdisciplinary reviews. Cognitive science, 7(5), 317–329.

Craig, A. D. (2002). How do you feel? Interoception: the sense of the physiological condition of the body. Nat Rev Neurosci, 3(8), 655–666.

Critchley, H. D., & Harrison, N. A. (2013). Visceral influences on brain and behavior. Neuron, 77(4), 624–638.

Critchley, H. D., Wiens, S., Rotshtein, P., Ohman, A., & Dolan, R. J. (2004). Neural systems supporting interoceptive awareness. Nat Neurosci, 7(2), 189–195.

Crittenden, B. M., Mitchell, D. J., & Duncan, J. (2016). Task Encoding across the Multiple Demand Cortex Is Consistent with a Frontoparietal and Cingulo-Opercular Dual Networks Distinction. The Journal of neuroscience : the official journal of the Society for Neuroscience, 36(23), 6147–6155.

Crockett, M. J., Braams, B. R., Clark, L., Tobler, P. N., Robbins, T. W., & Kalenscher, T. (2013). Restricting temptations: neural mechanisms of precommitment. Neuron, 79(2), 391–401.

Crone, E. A., Wendelken, C., Donohue, S. E., & Bunge, S. A. (2006). Neural evidence for dissociable components of task-switching. Cereb Cortex, 16(4), 475–486.

Crowe, D. A., Goodwin, S. J., Blackman, R. K., Sakellaridi, S., Sponheim, S. R., Macdonald, A. W., 3rd, et al. (2013). Prefrontal neurons transmit signals to parietal neurons that reflect executive control of cognition. Nat Neurosci, 16(10), 1484–1491.

Croxson, P. L., Walton, M. E., O’Reilly, J. X., Behrens, T. E., & Rushworth, M. F. (2009). Effort-based cost-benefit valuation and the human brain. J Neurosci, 29(14), 4531–4541.

D'Esposito, M., & Postle, B. R. (2015). The cognitive neuroscience of working memory. Annual Review of Psychology, 66, 115–142.

Davidson, R. J. (2000). Affective style, psychopathology, and resilience: brain mechanisms and plasticity. The American psychologist, 55(11), 1196–1214.

Daw, N. D., Niv, Y., & Dayan, P. (2005). Uncertainty-based competition between prefrontal and dorsolateral striatal systems for behavioral control. Nat Neurosci, 8(12), 1704–1711.

De Baene, W., Kuhn, S., & Brass, M. (2011). Challenging a decade of brain research on task switching: Brain activation in the task-switching paradigm reflects adaptation rather than reconfiguration of task sets. Hum Brain Mapp

Derrfuss, J., Brass, M., Neumann, J., & von Cramon, D. Y. (2005). Involvement of the inferior frontal junction in cognitive control: meta-analyses of switching and Stroop studies. Hum Brain Mapp, 25(1), 22–34.

Desimone, R., & Duncan, J. (1995). Neural mechanisms of selective visual attention. Annu Rev Neurosci, 18, 193–222.

Dias, R., Robbins, T. W., & Roberts, A. C. (1996). Dissociation in prefrontal cortex of affective and attentional shifts. Nature, 380(6569), 69–72.

Diekhof, E. K., & Gruber, O. (2010). When desire collides with reason: functional interactions between anteroventral prefrontal cortex and nucleus accumbens underlie the human ability to resist impulsive desires. J Neurosci, 30(4), 1488–1493.

Dixon, M. L. (2015). Cognitive Control, Emotional Value, and the Lateral Prefrontal Cortex. Frontiers in Psychology, 6.

Dixon, M. L., Andrews-Hanna, J. R., Spreng, R. N., Irving, Z. C., Mills, C., Girn, M., et al. (2017). Interactions between the default network and dorsal attention network vary across default subsystems, time, and cognitive states. Neuroimage, 147, 632–649.

Dixon, M. L., & Christoff, K. (2012). The Decision to Engage Cognitive Control is Driven by Expected Reward-Value: Neural and Behavioral Evidence. PLoS One, 7(12), 1–12.

Dixon, M. L., & Christoff, K. (2014). The lateral prefrontal cortex and complex value-based learning and decision making. Neurosci Biobehav Rev

Dixon, M. L., Fox, K. C. R., & Christoff, K. (2014a). Evidence for rostro-caudal functional organization in multiple brain areas related to goal-directed behavior. Brain Research

Dixon, M. L., Fox, K. C. R., & Christoff, K. (2014b). A framework for understanding the relationship between externally and internally directed cognition. Neuropsychologia, 62(321-330).

Dixon, M. L., Girn, M., & Christoff, K. (2017). Hierarchical organization of frontoparietal control networks underlying goal-directed behavior. In M. Watanabe (Ed.), The Prefrontal Cortex as an Executive, Emotional, and Social Brain (pp. 133–148): Springer Japan.

Dixon, M. L., Thiruchselvam, R., Todd, R., & Christoff, K. (2017). Emotion and the Prefrontal Cortex: An Integrative Review. Psychological bulletin.

Dosenbach, N. U., Fair, D. A., Miezin, F. M., Cohen, A. L., Wenger, K. K., Dosenbach, R. A., et al. (2007). Distinct brain networks for adaptive and stable task control in humans. Proc Natl Acad Sci U S A, 104(26), 11073–11078.

Dosenbach, N. U., Visscher, K. M., Palmer, E. D., Miezin, F. M., Wenger, K. K., Kang, H. C., et al. (2006). A core system for the implementation of task sets. Neuron, 50(5), 799–812.

Dumontheil, I., Thompson, R., & Duncan, J. (2011). Assembly and use of new task rules in fronto-parietal cortex. J Cogn Neurosci, 23(1), 168–182.

Duncan, J. (2001). An adaptive coding model of neural function in prefrontal cortex. Nat Rev Neurosci, 2(11), 820–829.

Duncan, J. (2010). The multiple-demand (MD) system of the primate brain: mental programs for intelligent behaviour. Trends Cogn Sci, 14(4), 172–179.

Duncan, J. (2013). The structure of cognition: attentional episodes in mind and brain. Neuron, 80(1), 35–50.

Egner, T., & Hirsch, J. (2005). Cognitive control mechanisms resolve conflict through cortical amplification of task-relevant information. Nat Neurosci, 8(12), 1784–1790.

Eickhoff, S. B., Bzdok, D., Laird, A. R., Kurth, F., & Fox, P. T. (2012). Activation likelihood estimation meta-analysis revisited. Neuroimage, 59(3), 2349–2361.

Eickhoff, S. B., Laird, A. R., Fox, P. M., Lancaster, J. L., & Fox, P. T. (2017). Implementation errors in the GingerALE Software: Description and recommendations. Human brain mapping, 38(1), 7–11.

Eickhoff, S. B., Laird, A. R., Grefkes, C., Wang, L. E., Zilles, K., & Fox, P. T. (2009). Coordinate-based activation likelihood estimation meta-analysis of neuroimaging data: a random-effects approach based on empirical estimates of spatial uncertainty. Human brain mapping, 30(9), 2907–2926.

Engelmann, J. B., Damaraju, E., Padmala, S., & Pessoa, L. (2009). Combined effects of attention and motivation on visual task performance: transient and sustained motivational effects. Front Hum Neurosci, 3, 4.

Essex, B. G., Clinton, S. A., Wonderley, L. R., & Zald, D. H. (2012). The Impact of the Posterior Parietal and Dorsolateral Prefrontal Cortices on the Optimization of Long-Term versus Immediate Value. J Neurosci, 32(44), 15403–15413.

Etzel, J. A., Cole, M. W., Zacks, J. M., Kay, K. N., & Braver, T. S. (2015). Reward Motivation Enhances Task Coding in Frontoparietal Cortex. Cereb Cortex.

Evans, A., Kamber, M., Collins, D., & MacDonald, D. (1994). An MRI-based probabilistic atlas of neuroanatomy Magnetic resonance scanning and epilepsy (pp. 263–274): Springer.

Farb, N. A., Segal, Z. V., & Anderson, A. K. (2012). Attentional Modulation of Primary Interoceptive and Exteroceptive Cortices. Cereb Cortex.

Fornito, A., Harrison, B. J., Zalesky, A., & Simons, J. S. (2012). Competitive and cooperative dynamics of large-scale brain functional networks supporting recollection. Proc Natl Acad Sci U S A, 109(31), 12788–12793.

Funahashi, S., Chafee, M. V., & Goldman-Rakic, P. S. (1993). Prefrontal neuronal activity in rhesus monkeys performing a delayed anti-saccade task. Nature, 365(6448), 753–756.

Fuster, J. M. (1989). The prefrontal cortex : anatomy, physiology, and neuropsychology of the frontal lobe (2nd ed.). New York: Raven Press.

Fuster, J. M., & Alexander, G. E. (1971). Neuron activity related to short-term memory. Science, 173(3997), 652–654.

Gao, W., & Lin, W. (2012). Frontal parietal control network regulates the anti-correlated default and dorsal attention networks. Hum Brain Mapp, 33(1), 192–202.

Gerlach, K. D., Spreng, R. N., Madore, K. P., & Schacter, D. L. (2014). Future planning: Default network activity couples with frontoparietal control network and reward-processing regions during process and outcome simulations. Soc Cogn Affect Neurosci.

Gianotti, L. R., Knoch, D., Faber, P. L., Lehmann, D., Pascual-Marqui, R. D., Diezi, C., et al. (2009). Tonic activity level in the right prefrontal cortex predicts individuals' risk taking. Psychol Sci, 20(1), 33–38.

Gilbert, A. M., & Fiez, J. A. (2004). Integrating rewards and cognition in the frontal cortex. Cognitive, affective & behavioral neuroscience, 4(4), 540–552.

Goldman-Rakic, P. S. (1987). Circuitry of primate prefrontal cortex and regulation of behavior by representational memory. In F. Plum (Ed.), Handbook of Physiology: The Nervous System (pp. 373–417). Bethesda, MD: American Physiological Society.

Gollwitzer, P. M. (1999). Implementation intentions: strong effects of simple plans. American Psychologist, 54(7), 493–503.

Hare, T. A., O'Doherty, J., Camerer, C. F., Schultz, W., & Rangel, A. (2008). Dissociating the role of the orbitofrontal cortex and the striatum in the computation of goal values and prediction errors. J Neurosci, 28(22), 5623–5630.

Hazy, T. E., Frank, M. J., & O'Reilly R, C. (2007). Towards an executive without a homunculus: computational models of the prefrontal cortex/basal ganglia system. Philosophical transactions of the Royal Society of London. Series B, Biological sciences, 362(1485), 1601–1613.

Hazy, T. E., Frank, M. J., & O'Reilly, R. C. (2006). Banishing the homunculus: making working memory work. Neuroscience, 139(1), 105–118.

Heller, A. S., Johnstone, T., Shackman, A. J., Light, S. N., Peterson, M. J., Kolden, G. G., et al. (2009). Reduced capacity to sustain positive emotion in major depression reflects diminished maintenance of fronto-striatal brain activation. Proc Natl Acad Sci U S A, 106(52), 22445–22450.

Hikosaka, K., & Watanabe, M. (2000). Delay activity of orbital and lateral prefrontal neurons of the monkey varying with different rewards. Cereb Cortex, 10(3), 263–271.

Histed, M. H., Pasupathy, A., & Miller, E. K. (2009). Learning substrates in the primate prefrontal cortex and striatum: sustained activity related to successful actions. Neuron, 63(2), 244–253.

Hosokawa, T., & Watanabe, M. (2012). Prefrontal Neurons Represent Winning and Losing during Competitive Video Shooting Games between Monkeys. J Neurosci, 32(22), 7662–7671.

Huettel, S. A., Song, A. W., & McCarthy, G. (2005). Decisions under uncertainty: probabilistic context influences activation of prefrontal and parietal cortices. J Neurosci, 25(13), 3304–3311.

Hutcherson, C. A., Plassmann, H., Gross, J. J., & Rangel, A. (2012). Cognitive regulation during decision making shifts behavioral control between ventromedial and dorsolateral prefrontal value systems. J Neurosci, 32(39), 13543–13554.

Ivanov, I., Liu, X., Clerkin, S., Schulz, K., Friston, K., Newcorn, J. H., et al. (2012). Effects of motivation on reward and attentional networks: an fMRI study. Brain and Behavior, 2(6), 741–753.

Jezzini, A., Caruana, F., Stoianov, I., Gallese, V., & Rizzolatti, G. (2012). Functional organization of the insula and inner perisylvian regions. Proc Natl Acad Sci U S A, 109(25), 10077–10082.

Jimura, K., Chushak, M. S., & Braver, T. S. (2013). Impulsivity and self-control during intertemporal decision making linked to the neural dynamics of reward value representation. J Neurosci, 33(1), 344–357.

Jimura, K., Locke, H. S., & Braver, T. S. (2010). Prefrontal cortex mediation of cognitive enhancement in rewarding motivational contexts. Proc Natl Acad Sci U S A, 107(19), 8871–8876.

Jimura, K., & Poldrack, R. A. (2012). Analyses of regional-average activation and multivoxel pattern information tell complementary stories. Neuropsychologia, 50(4), 544–552.

Kaiser, R. H., Andrews-Hanna, J. R., Spielberg, J. M., Warren, S. L., Sutton, B. P., Miller, G. A., et al. (2015). Distracted and down: neural mechanisms of affective interference in subclinical depression. Social cognitive and affective neuroscience, 10(5), 654–663.

Kaiser, R. H., Andrews-Hanna, J. R., Wager, T. D., & Pizzagalli, D. A. (2015). Large-Scale Network Dysfunction in Major Depressive Disorder: A Meta-analysis of Resting-State Functional Connectivity. JAMA Psychiatry, 72(6), 603–611.

Kennerley, S. W., & Wallis, J. D. (2009). Evaluating choices by single neurons in the frontal lobe: outcome value encoded across multiple decision variables. Eur J Neurosci, 29(10), 2061–2073.

Kim, S., Hwang, J., & Lee, D. (2008). Prefrontal coding of temporally discounted values during intertemporal choice. Neuron, 59(1), 161–172.

Klein, J. T., Deaner, R. O., & Platt, M. L. (2008). Neural correlates of social target value in macaque parietal cortex. Current biology : CB, 18(6), 419–424.

Knutson, B., Adams, C. M., Fong, G. W., & Hommer, D. (2001). Anticipation of increasing monetary reward selectively recruits nucleus accumbens. J Neurosci, 21(16), RC159.

Knutson, B., & Greer, S. M. (2008). Anticipatory affect: neural correlates and consequences for choice. Philos Trans R Soc Lond B Biol Sci, 363(1511), 3771–3786.

Kobayashi, S., Nomoto, K., Watanabe, M., Hikosaka, O., Schultz, W., & Sakagami, M. (2006). Influences of rewarding and aversive outcomes on activity in macaque lateral prefrontal cortex. Neuron, 51(6), 861–870.

Koechlin, E., Ody, C., & Kouneiher, F. (2003). The architecture of cognitive control in the human prefrontal cortex. Science, 302(5648), 1181–1185.

Kool, W., McGuire, J. T., Rosen, Z. B., & Botvinick, M. M. (2010). Decision making and the avoidance of cognitive demand. J Exp Psychol Gen, 139(4), 665–682.

Kouneiher, F., Charron, S., & Koechlin, E. (2009). Motivation and cognitive control in the human prefrontal cortex. Nat Neurosci, 12(7), 939–945.

Krebs, R. M., Boehler, C. N., Roberts, K. C., Song, A. W., & Woldorff, M. G. (2012). The involvement of the dopaminergic midbrain and cortico-striatal-thalamic circuits in the integration of reward prospect and attentional task demands. Cerebral cortex, 22(3), 607–615.

Kurniawan, I. T., Guitart-Masip, M., Dayan, P., & Dolan, R. J. (2013). Effort and valuation in the brain: the effects of anticipation and execution. J Neurosci, 33(14), 6160–6169.

Laird, A. R., Fox, P. M., Price, C. J., Glahn, D. C., Uecker, A. M., Lancaster, J. L., et al. (2005). ALE meta-analysis: controlling the false discovery rate and performing statistical contrasts. Human brain mapping, 25(1), 155–164.

Lebreton, M., Bertoux, M., Boutet, C., Lehericy, S., Dubois, B., Fossati, P., et al. (2013). A critical role for the hippocampus in the valuation of imagined outcomes. PLoS Biol, 11(10), e1001684.

Leon, M. I., & Shadlen, M. N. (1999). Effect of expected reward magnitude on the response of neurons in the dorsolateral prefrontal cortex of the macaque. Neuron, 24(2), 415–425.

Leotti, L. A., & Delgado, M. R. (2014). The value of exercising control over monetary gains and losses. Psychol Sci, 25(2), 596–604.

Levy, D. J., & Glimcher, P. W. (2012). The root of all value: a neural common currency for choice. Current opinion in neurobiology, 22(6), 1027–1038.

Locke, H. S., & Braver, T. S. (2008). Motivational influences on cognitive control: behavior, brain activation, and individual differences. Cogn Affect Behav Neurosci, 8(1), 99–112.

Matsumoto, K., Suzuki, W., & Tanaka, K. (2003). Neuronal correlates of goal-based motor selection in the prefrontal cortex. Science, 301(5630), 229–232.

McClure, S. M., Laibson, D. I., Loewenstein, G., & Cohen, J. D. (2004). Separate neural systems value immediate and delayed monetary rewards. Science, 306(5695), 503–507.

McGuire, J. T., & Botvinick, M. M. (2010). Prefrontal cortex, cognitive control, and the registration of decision costs. Proc Natl Acad Sci U S A, 107(17), 7922–7926.

Medford, N., & Critchley, H. D. (2010). Conjoint activity of anterior insular and anterior cingulate cortex: awareness and response. Brain Struct Funct, 214(5-6), 535–549.

Meiran, N. (1996). Reconfiguration of processing mode prior to task performance. Journal of Experimental Psychology: Learning, Memory, and Cognition, 22(6), 1423.

Meiran, N. (2000). Modeling cognitive control in task-switching. Psychological research, 63(3-4), 234–249.

Menon, V., & Uddin, L. Q. (2010). Saliency, switching, attention and control: a network model of insula function. Brain Struct Funct, 214(5-6), 655–667.

Miller, E. K., & Cohen, J. D. (2001). An integrative theory of prefrontal cortex function. Annu Rev Neurosci, 24, 167–202.

Mitchell, D. J., Bell, A. H., Buckley, M. J., Mitchell, A. S., Sallet, J., & Duncan, J. (2016). A Putative Multiple-Demand System in the Macaque Brain. The Journal of neuroscience : the official journal of the Society for Neuroscience, 36(33), 8574–8585.

Miyake, A., Friedman, N. P., Emerson, M. J., Witzki, A. H., Howerter, A., & Wager, T. D. (2000). The unity and diversity of executive functions and their contributions to complex “Frontal Lobe” tasks: a latent variable analysis. Cognitive psychology, 41(1), 49–100.

Montague, P. R., Dayan, P., & Sejnowski, T. J. (1996). A framework for mesencephalic dopamine systems based on predictive Hebbian learning. The Journal of neuroscience : the official journal of the Society for Neuroscience, 16(5), 1936–1947.

Morecraft, R. J., & Tanji, J. (2009). Cingulofrontal interactions and the cingulate motor areas. In B. A. Vogt (Ed.), Cingulate Neurobiology and Disease (pp. 113–144). Oxford, New York: Oxford University Press.

Munakata, Y., Herd, S. A., Chatham, C. H., Depue, B. E., Banich, M. T., & O'Reilly, R. C. (2011). A unified framework for inhibitory control. Trends Cogn Sci, 15(10), 453–459.

Nigg, J. T., & Casey, B. J. (2005). An integrative theory of attention-deficit/ hyperactivity disorder based on the cognitive and affective neurosciences. Dev Psychopathol, 17(3), 785–806.

O'Doherty, J., Dayan, P., Schultz, J., Deichmann, R., Friston, K., & Dolan, R. J. (2004). Dissociable roles of ventral and dorsal striatum in instrumental conditioning. Science, 304(5669), 452–454.

O'Doherty, J. P. (2004). Reward representations and reward-related learning in the human brain: insights from neuroimaging. Curr Opin Neurobiol, 14(6), 769–776.

O'Reilly, R. C., Herd, S. A., & Pauli, W. M. (2010). Computational models of cognitive control. Current opinion in neurobiology, 20(2), 257–261.

Padmala, S., & Pessoa, L. (2011). Reward reduces conflict by enhancing attentional control and biasing visual cortical processing. J Cogn Neurosci, 23(11), 3419–3432.

Pan, X., Sawa, K., Tsuda, I., Tsukada, M., & Sakagami, M. (2008). Reward prediction based on stimulus categorization in primate lateral prefrontal cortex. Nat Neurosci, 11(6), 703–712.

Paradiso, S., Chemerinski, E., Yazici, K. M., Tartaro, A., & Robinson, R. G. (1999). Frontal lobe syndrome reassessed: comparison of patients with lateral or medial frontal brain damage. J Neurol Neurosurg Psychiatry, 67(5), 664–667.

Parvizi, J., Rangarajan, V., Shirer, W. R., Desai, N., & Greicius, M. D. (2013). The will to persevere induced by electrical stimulation of the human cingulate gyrus. Neuron, 80(6), 1359–1367.

Paschke, L. M., Walter, H., Steimke, R., Ludwig, V. U., Gaschler, R., Schubert, T., et al. (2015). Motivation by potential gains and losses affects control processes via different mechanisms in the attentional network. Neuroimage, 111, 549–561.

Passingham, R. E., & Wise, S. P. (2012). The neurobiology of the prefrontal cortex : anatomy, evolution, and the origin of insight (1st ed.). Oxford: Oxford University Press.

Pessoa, L. (2008). On the relationship between emotion and cognition. Nat Rev Neurosci, 9(2), 148–158.

Plassmann, H., O'Doherty, J., & Rangel, A. (2007). Orbitofrontal cortex encodes willingness to pay in everyday economic transactions. J Neurosci, 27(37), 9984–9988.

Plassmann, H., O'Doherty, J. P., & Rangel, A. (2010). Appetitive and aversive goal values are encoded in the medial orbitofrontal cortex at the time of decision making. J Neurosci, 30(32), 10799–10808.

Platt, M. L., & Glimcher, P. W. (1999). Neural correlates of decision variables in parietal cortex. Nature, 400(6741), 233–238.

Pochon, J. B., Levy, R., Fossati, P., Lehericy, S., Poline, J. B., Pillon, B., et al. (2002). The neural system that bridges reward and cognition in humans: an fMRI study. Proc Natl Acad Sci U S A, 99(8), 5669–5674.

Posner, M. I., & Dehaene, S. (1994). Attentional networks. Trends Neurosci, 17(2), 75–79.

Posner, M. I., & DiGirolamo, G. J. (1998). Conflict, target detection and cognitive control. The attentive brain, 401–423.

Power, J. D., Cohen, A. L., Nelson, S. M., Wig, G. S., Barnes, K. A., Church, J. A., et al. (2011). Functional network organization of the human brain. Neuron, 72(4), 665–678.

Procyk, E., Wilson, C. R., Stoll, F. M., Faraut, M. C., Petrides, M., & Amiez, C. (2014). Midcingulate Motor Map and Feedback Detection: Converging Data from Humans and Monkeys. Cereb Cortex

Rangel, A., Camerer, C., & Montague, P. R. (2008). A framework for studying the neurobiology of value-based decision making. Nat Rev Neurosci, 9(7), 545–556.

Rangel, A., & Hare, T. (2010). Neural computations associated with goal-directed choice. Curr Opin Neurobiol, 20(2), 262–270.

Ridderinkhof, K. R., Ullsperger, M., Crone, E. A., & Nieuwenhuis, S. (2004). The role of the medial frontal cortex in cognitive control. Science, 306(5695), 443–447.

Rorden, C., Karnath, H. O., & Bonilha, L. (2007). Improving lesion-symptom mapping. Journal of Cognitive Neuroscience, 19(7), 1081–1088.

Rowe, J. B., Eckstein, D., Braver, T., & Owen, A. M. (2008). How does reward expectation influence cognition in the human brain? Journal of Cognitive Neuroscience, 20(11), 1980–1992.

Rushworth, M. F., Behrens, T. E., Rudebeck, P. H., & Walton, M. E. (2007). Contrasting roles for cingulate and orbitofrontal cortex in decisions and social behaviour. Trends Cogn Sci, 11(4), 168–176.

Rushworth, M. F., Noonan, M. P., Boorman, E. D., Walton, M. E., & Behrens, T. E. (2011). Frontal cortex and reward-guided learning and decision-making. Neuron, 70(6), 1054–1069.

Rushworth, M. F., Passingham, R. E., & Nobre, A. C. (2002). Components of switching intentional set. J Cogn Neurosci, 14(8), 1139–1150.

Schoenbaum, G., & Esber, G. R. (2010). How do you (estimate you will) like them apples? Integration as a defining trait of orbitofrontal function. Curr Opin Neurobiol, 20(2), 205–211.

Schultz, W. (1997). Dopamine neurons and their role in reward mechanisms. Curr Opin Neurobiol, 7(2), 191–197.

Seeley, W. W., Menon, V., Schatzberg, A. F., Keller, J., Glover, G. H., Kenna, H., et al. (2007). Dissociable intrinsic connectivity networks for salience processing and executive control. J Neurosci, 27(9), 2349–2356.

Seo, H., Barraclough, D. J., & Lee, D. (2007). Dynamic signals related to choices and outcomes in the dorsolateral prefrontal cortex. Cereb Cortex, 17 Suppl 1, i110–117.

Shackman, A. J., Salomons, T. V., Slagter, H. A., Fox, A. S., Winter, J. J., & Davidson, R. J. (2011). The integration of negative affect, pain and cognitive control in the cingulate cortex. Nat Rev Neurosci, 12(3), 154–167.

Shackman, A. J., Tromp, D. P., Stockbridge, M. D., Kaplan, C. M., Tillman, R. M., & Fox, A. S. (2016). Dispositional negativity: An integrative psychological and neurobiological perspective. Psychological bulletin, 142(12), 1275–1314.

Shenhav, A., Botvinick, M. M., & Cohen, J. D. (2013). The expected value of control: an integrative theory of anterior cingulate cortex function. Neuron, 79(2), 217–240.

Shidara, M., & Richmond, B. J. (2002). Anterior cingulate: single neuronal signals related to degree of reward expectancy. Science, 296(5573), 1709–1711.

Shima, K., & Tanji, J. (1998). Role for cingulate motor area cells in voluntary movement selection based on reward. Science, 282(5392), 1335–1338.

Soutschek, A., Stelzel, C., Paschke, L., Walter, H., & Schubert, T. (2015). Dissociable effects of motivation and expectancy on conflict processing: an fMRI study. Journal of Cognitive Neuroscience, 27(2), 409–423.

Spreng, R. N., Stevens, W. D., Chamberlain, J. P., Gilmore, A. W., & Schacter, D. L. (2010). Default network activity, coupled with the frontoparietal control network, supports goal-directed cognition. Neuroimage, 53(1), 303–317.

Stokes, M. G., Kusunoki, M., Sigala, N., Nili, H., Gaffan, D., & Duncan, J. (2013). Dynamic coding for cognitive control in prefrontal cortex. Neuron, 78(2), 364–375.

Stuss, D. T., & Knight, R. T. (2002). Principles of frontal lobe function. Oxford; New York: Oxford University Press.

Tanaka, S. C., Doya, K., Okada, G., Ueda, K., Okamoto, Y., & Yamawaki, S. (2004). Prediction of immediate and future rewards differentially recruits cortico-basal ganglia loops. Nat Neurosci, 7(8), 887–893.

Taylor, S. F., Welsh, R. C., Wager, T. D., Phan, K. L., Fitzgerald, K. D., & Gehring, W. J. (2004). A functional neuroimaging study of motivation and executive function. Neuroimage, 21(3), 1045–1054.

Tobler, P. N., Christopoulos, G. I., O'Doherty, J. P., Dolan, R. J., & Schultz, W. (2009). Risk-dependent reward value signal in human prefrontal cortex. Proc Natl Acad Sci U S A, 106(17), 7185–7190.

Tomita, H., Ohbayashi, M., Nakahara, K., Hasegawa, I., & Miyashita, Y. (1999). Top-down signal from prefrontal cortex in executive control of memory retrieval. Nature, 401(6754), 699–703.

Turkeltaub, P. E., Eickhoff, S. B., Laird, A. R., Fox, M., Wiener, M., & Fox, P. (2012). Minimizing within-experiment and within-group effects in Activation Likelihood Estimation meta-analyses. Human brain mapping, 33(1), 1–13.

Ullsperger, M., Danielmeier, C., & Jocham, G. (2014). Neurophysiology of performance monitoring and adaptive behavior. Physiological reviews, 94(1), 35–79.

Vickery, T. J., Chun, M. M., & Lee, D. (2011). Ubiquity and specificity of reinforcement signals throughout the human brain. Neuron, 72(1), 166–177.

Vincent, J. L., Kahn, I., Snyder, A. Z., Raichle, M. E., & Buckner, R. L. (2008). Evidence for a frontoparietal control system revealed by intrinsic functional connectivity. J Neurophysiol, 100(6), 3328–3342.

Wallis, J. D., Anderson, K. C., & Miller, E. K. (2001). Single neurons in prefrontal cortex encode abstract rules. Nature, 411(6840), 953–956.

Wallis, J. D., & Miller, E. K. (2003). Neuronal activity in primate dorsolateral and orbital prefrontal cortex during performance of a reward preference task. Eur J Neurosci, 18(7), 2069–2081.

Watanabe, M. (1996). Reward expectancy in primate prefrontal neurons. Nature, 382(6592), 629–632.

Watanabe, M. (2017). The Prefrontal Cortex as an Executive, Emotional, and Social Brain: Springer.

Watanabe, M., Hikosaka, K., Sakagami, M., & Shirakawa, S. (2002). Coding and monitoring of motivational context in the primate prefrontal cortex. J Neurosci, 22(6), 2391–2400.

Watanabe, M., & Sakagami, M. (2007). Integration of cognitive and motivational context information in the primate prefrontal cortex. Cereb Cortex, 17 Suppl 1, i101–109.

Weber, B. J., & Huettel, S. A. (2008). The neural substrates of probabilistic and intertemporal decision making. Brain Res, 1234, 104–115.

Westbrook, A., Kester, D., & Braver, T. S. (2013). What is the subjective cost of cognitive effort? Load, trait, and aging effects revealed by economic preference. PLoS One, 8(7), e68210.

Yarkoni, T., Poldrack, R. A., Nichols, T. E., Van Essen, D. C., & Wager, T. D. (2011). Large-scale automated synthesis of human functional neuroimaging data. Nat Methods, 8(8), 665–670.

Yeo, B. T., Krienen, F. M., Sepulcre, J., Sabuncu, M. R., Lashkari, D., Hollinshead, M., et al. (2011). The organization of the human cerebral cortex estimated by intrinsic functional connectivity. J Neurophysiol, 106(3), 1125–1165.

Zamboni, G., Huey, E. D., Krueger, F., Nichelli, P. F., & Grafman, J. (2008). Apathy and disinhibition in frontotemporal dementia: Insights into their neural correlates. Neurology, 71(10), 736–742.

